# Development of a latency model based on HIV-1 subtype C to study how long terminal repeat genetic variation impacts viral persistence and latency reversal

**DOI:** 10.1101/2024.08.02.606371

**Authors:** Shreyal Maikoo, Robert-Jan Palstra, Krista L. Dong, Tokameh Mahmoudi, Thumbi Ndung’u, Paradise Madlala

## Abstract

Most people living with HIV (PLWH) reside in sub-Saharan Africa. South Africa is the epicentre where 98% of HIV-1 infections are subtype C. However, partially due to unavailability of non-subtype B latency models, most studies of HIV-1 latency and cure have focused on HIV-1 subtype B (HIV-1B) which predominates Europe and USA. Moreover, the effect of inter- and intra-subtype genetic variation of the viral promoter, long terminal repeat (LTR), from PLWH on latency reversal is unknown. We constructed a retroviral vector expressing green fluorescent protein and HIV-1 subtype C (HIV-1C) consensus Tat protein under the control of either HIV-1C consensus or PLWH-derived transmitted/founder (T/F) LTR, produced respective LTR pseudotyped viruses, infected Jurkat E6 and primary CD4+ T cells *in vitro*, enriched for latently infected cells, and treated these cells with different latency reversing agents. We show that the HIV-1C LTR exhibited lower reactivation compared to HIV-1B. Furthermore, HIV-1C T/F LTR pseudotyped proviral variants with four NF-κB motifs exhibited lower reactivation compared to those with three NF-κB motifs. Our data indicate that inter- and intra-subtype HIV-1 LTR genetic variation in combination with host variation modulates latency reversal.

**Author summary:** Antiretroviral therapy (ART) suppresses HIV-1 replication, but it is not curative due to a persistent latent reservoir established early in infection. Although HIV-1 subtype C (HIV-1C) is responsible for about 50% of global and 98% of southern Africa infections, it is underrepresented in HIV-1 cure studies. The unavailability of non-subtype B latency models has led to most studies on HIV-1 latency and cure focusing on HIV-1 subtype B (HIV-1B) which predominates in western countries. The viral promoter, long terminal repeat (LTR) drives viral gene transcription and is important for the HIV-1 life cycle. In this study we undertook to develop a latency model based on HIV-1 subtype C to investigate the effect of inter- and intra- LTR genetic variation on viral persistence and latency reversal. Our data show that the HIV-1C LTR exhibited lower reactivation compared to HIV-1B. Furthermore, HIV-1C T/F LTR pseudotyped proviral variants with four NF-κB motifs exhibited lower reactivation compared to those with three NF-κB motifs. Taken together, our data suggest that inter- and intra-subtype HIV-1 LTR genetic variation in combination with host variation modulates latency reversal.

## Introduction

The persistence of the latent viral reservoir despite active antiretroviral therapy (ART) is the major barrier to HIV-1 infection cure [1]. The latent reservoir comprises of cells infected with replication competent but transcriptionally silent proviruses [2, 3]. One strategy being pursued to clear latently infected cells is to stimulate virus production from latent provirus using latency reversing agents (LRAs) (reviewed in [4]).

Different classes of LRAs include histone deacetylase inhibitors (HDACIs) [5, 6], protein kinase C (PKC) agonists [7, 8], and inhibitors of BAF (BAFi). HDACIs hyperacetylate histones resulting in less condensed chromatin thus allowing for host transcription factors to bind the integrated proviral promoter, i.e. the 5’ long terminal repeat (LTR), and induce viral gene expression [9]. The PKC agonists such as phorbol 12-myristate 13-acetate (PMA) plus Ionomycin (IONO) and TNF-α reactivate viral gene transcription through NF-ĸB signaling (Castro-Gonzalez et al., 2018). However, clinical trial results are not promising with different LRAs inducing differential reactivation and no reduction in the viral reservoir in people living with HIV-1 (PLWH) who are on antiretroviral therapy (ART) [5, 6, 9–11].

The HIV-1 5’ LTR is the viral promoter that drives viral gene transcription [12]. The HIV-1 5’ LTR is identical to the 3’ LTR, and is divided into three distinct regions, U3, R and U5. Specifically, the U3 region is further divided into three functional domains that regulate HIV-1 positive sense transcription: a core promoter region, enhancer region and modulatory region ((reviewed in [13]). HIV-1 subtype AE has only one NF-κB motif within the enhancer region, but most subtypes including the prototype subtype B have two, while subtype C has three to four NF-κB motifs that may translate to functional differences [14, 15]. The HIV-1 subtype C (HIV-1C) strains exhibiting LTR variants with four NF-κB motifs circulate at a low frequency in Brazil, Mozambique [16] and South Africa [17, 18]. The HIV-1C viral strains exhibiting 4 NF-κB motifs were reported to be taking over the epidemic in India and were associated with higher transcription activity and viral load compared to the standard subtype C LTR variants exhibiting three NF-κB motifs [19]. Using a recent-infection FRESH cohort established in Durban, South Africa in 2012, we recently confirmed that HIV-1C LTR variants exhibiting four NF-κB motifs are infrequent in South Africa and that genetic variation of HIV-1C transmitted/founder (T/F) LTR impacts transcription activation and clinical disease outcomes in ART-naïve PLWH [20].

However, the role of inter- and intra-subtype LTR genetic variation on the propensity of latency reversal has not been fully investigated. Therefore, in this study we undertook to develop a subtype C-based *in vitro* latency model, referred to as the C J-Lat cell line by adapting methods used to develop the J-Lat model for subtype B [21] and characterised the effect of HIV-1 LTR genetic variation on latency reactivation potential. We report the successful construction of an HIV-1C lentiviral minimal genome reporter vector expressing a green fluorescent protein (GFP) reporter and HIV-1C consensus Tat under the control of an HIV-1C consensus LTR (subtype C LTR-Tat-IRES-EGFP), referred to as C731CC and the development of a subtype C *in vitro* latency model (C J-Lat). Specifically, we show that HIV-1C is significantly less sensitive to reactivation by different LRAs compared to HIV-1B and HIV-1C T/F LTR variants display differential reactivation with four NF-κB LTR variants generally exhibiting lower propensity of latency reversal compared to the standard subtype C LTR in latently infected Jurkat E6 and primary CD4^+^ T-cells.

## Results

### Successful development of C J-Lat model

Genetic variation within an AP-1 [22] and the higher number of NF-κB binding sites [23] were reported to be associated with rapid latency establishment and a stable viral latent state. On the other hand, a different study reported no gross differences among subtype specific LTR variants, both in the initial latency level and the activation response, except for subtype AE [24]. A recent study reported that genetic variation of women living with HIV-1C derived LTR influence the establishment of latency [25]. However, the effect of inter-subtype and PLWH derived LTR genetic variation on the propensity of latency reactivation has not been fully investigated. Therefore, we hypothesized that inter- and intra-subtype LTR genetic variation may mediate latency reversal.

First, we successfully developed a C J-Lat model. The C731CC viruses we generated (see methods) were infectious resulting in up to 99% of GFP positive cells (Fig 1A). Virus dilutions exhibiting approximately 5% GFP positive cells were used to enrich for latently infected cells on the fourth day following the infection and overnight reactivation with different LRAs (Fig 1A). The concentrations of LRAs used in this study were not toxic to the cells, with the cell viability of PMA-treated cells being 95% as a representative. Following reactivation with PMA, TNF-α, prostratin and SAHA we found that 0.38%; 0.26%; 0.11%; and 0.04% of Jurkat E6 cells latently infected with C731CC proviruses expressed GFP compared to 0.68%; 0.5%; 0.2%; and 0.11% GFP expressing latently B731BB provirus infected cells (Fig 1B). Western blot analysis demonstrated that the expression levels of B731BB Tat (Tat B) were similar to C731CC provirus Tat (Tat C) levels in Jurkat E6 (Fig 1C). Equally, there were no differences in the amount of integrated HIV-1 DNA copies between B731BB and C731CC (Fig 1D). The main differences between the B- and C-subtype LTRs are differences in the number of NF-kB motifs, with B731BB proviruses exhibiting two and C731CC exhibiting three NF-κB motifs. In addition, the B731BB proviruses contain a four-nucleotide AP-1 sequence (TGAC), whereas the C731CC proviruses contain a seven-nucleotide AP-1 sequence (TGACACA) (Fig S1). Taken together, our data suggest that HIV-1C is two-fold less sensitive to reactivation with different LRAs compared to HIV-1B.

**Fig 1:**
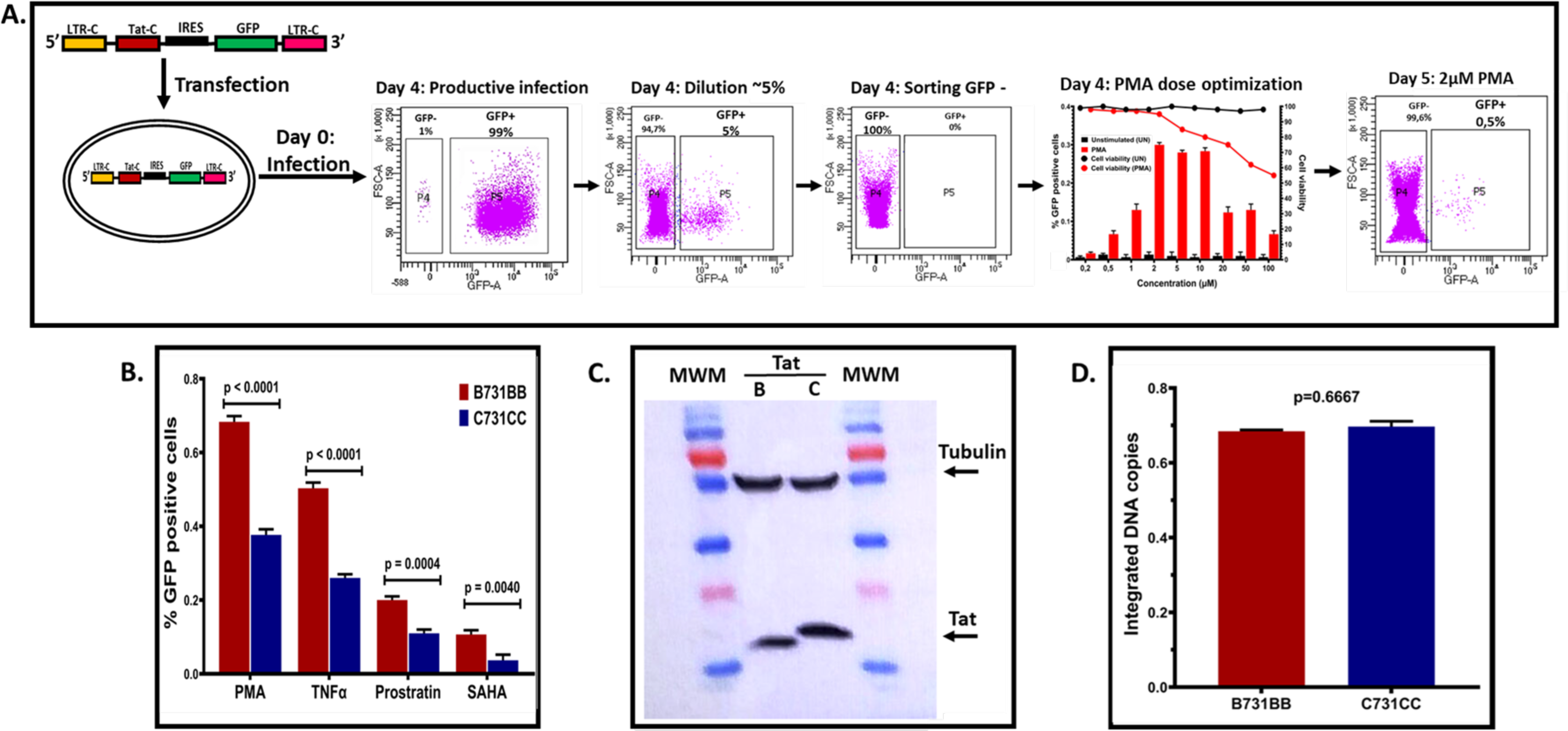
Outline of latency establishment experiment and development of C J-Lat model. **A:** Overview of transfection of the C731CC lentiviral vector into the Jurkat cells which resulted in productive infection (undiluted virus), as well as the dilution that gave ∼5% GFP positive cells. We then sorted for the remaining 95% GFP negative population (which gave 0% GFP positive cells) and tested a range of PMA concentrations to determine the optimal dose with least toxicity to the cells. 2 µM PMA was then selected as the optimal concentration, which was then used to reactivate all latent viruses. **B:** B731BB (subtype B-red bar) was twice as more reactivatable as compared to C731CC (subtype C-blue bar) when stimulated with LRAs. **C:** Western blot demonstrating that HIV-1 Tat was equally expressed in both subtypes (Tat C is HIV-1 subtype C Tat, whilst Tat B is HIV-1 subtype B Tat). Tubulin was used as a loading control. **D:** Alu-gag PCR demonstrating that there was no significant difference in the amount of integrated HIV-1 DNA copies between B731BB and C731CC J-Lat lines.

### Phylogenetic analysis of participant -derived HIV-1C T/F LTRs

We recently showed that genetic variation of the HIV-1 subtype C T/F LTR impacts transcription activation potential and clinical disease outcomes in ART-naïve PLWH [20]. However, the effect of T/F LTR genetic variation on the propensity of latency reactivation remains unknown. We hypothesized that PLWH derived HIV-1C T/F LTR variants modulate the propensity for latency reversal. To this effect, we replaced the consensus LTR of the C731CC lentiviral vector with individual HIV-1C T/F LTR variants. Phylogenetic analysis confirmed that PLWH derived T/F LTR sequences belonged to HIV-1C as they rooted to South African HIV-1C reference sequence and that they were unlinked since they did not cluster together (Fig 2A). Moreover, the T/F LTR sequences cloned into C731CC had the same branch length and clustered together with the corresponding T/F LTR sequences contained in pGL3 published in our previous study [20], We randomly selected 20 T/F LTR sequences to produce pseudotyped viruses which are highlighted in blue text (Fig 2A). The T/F LTR pseudotyped viruses were used to infect the Jurkat cells (Fig 2B) and GFP negative cells were sorted from a virus dilution that yielded approximately 5% GFP positive cells.

**Fig 2:**
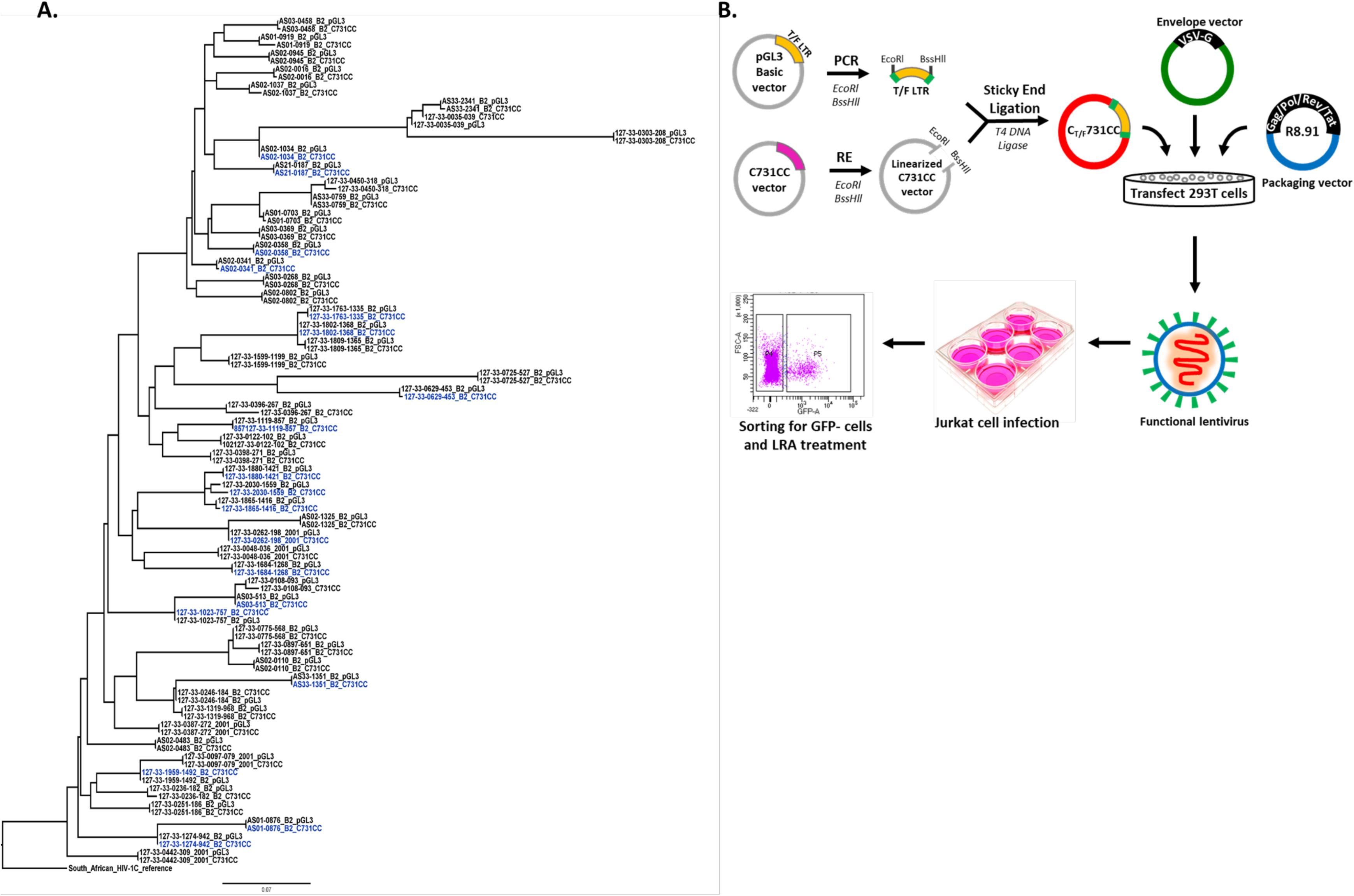
Phylogenetic analysis of participant derived HIV-1C T/F LTR sequences before and after cloning confirms successful cloning. **A:** Phylogenetic tree that was constructed using the online tool Phyml (http://www.hivlanl.gov) and rerooted on the South African subtype C reference sequence using Figtree software v1.4.3. The HIV-1C T/F LTR sequences obtained before cloning are denoted as pGL3, whilst those obtained after cloning are denoted as C731CC. The 20 LTR sequences highlighted in blue were randomly selected to produce the participant -derived HIV-1C T/F LTR pseudotyped viruses. The HIV-1C LTR for each participant identity (PID) from both pGL3 and C731CC clusters together thus demonstrating that the HIV-1C LTR sequence was the same before and after cloning, suggesting that cloning was successful with no sequence recombination. **B:** Overview of the methodology undertaken to produce the participant -derived HIV-1C T/F LTR pseudotyped viruses.

### Effect of participant -derived HIV-1C T/F LTR on latency reactivation

Our data demonstrate that all 20 participant-derived HIV-1C T/F LTR pseudotyped viruses (denoted as Pt 1-20 henceforth) exhibited differential reactivation by all four different LRAs (heatmap in Fig 3 and raw data in Fig S2 and Table S1). Unsupervised clustering generated five clusters: two clusters (Pt 2 & 15 and Pt 1, 8, & 9) showing overall higher reactivation; one cluster (Pt 3, 11, 14 & 16) showing moderately higher or mixed reactivation; one cluster (Pt 6, 10, 17, 18, 19, 20) showing moderately lower reactivation and the last cluster (Pt 4, 5, 7, 12 &13) showing overall lower reactivation (Fig 3) as compared to the average reactivation for all LRAs. This suggests that PLWH derived HIV-1C T/F LTR variants exhibit differential sensitivity to LRAs and thus have different reactivation potential.

**Fig 3:**
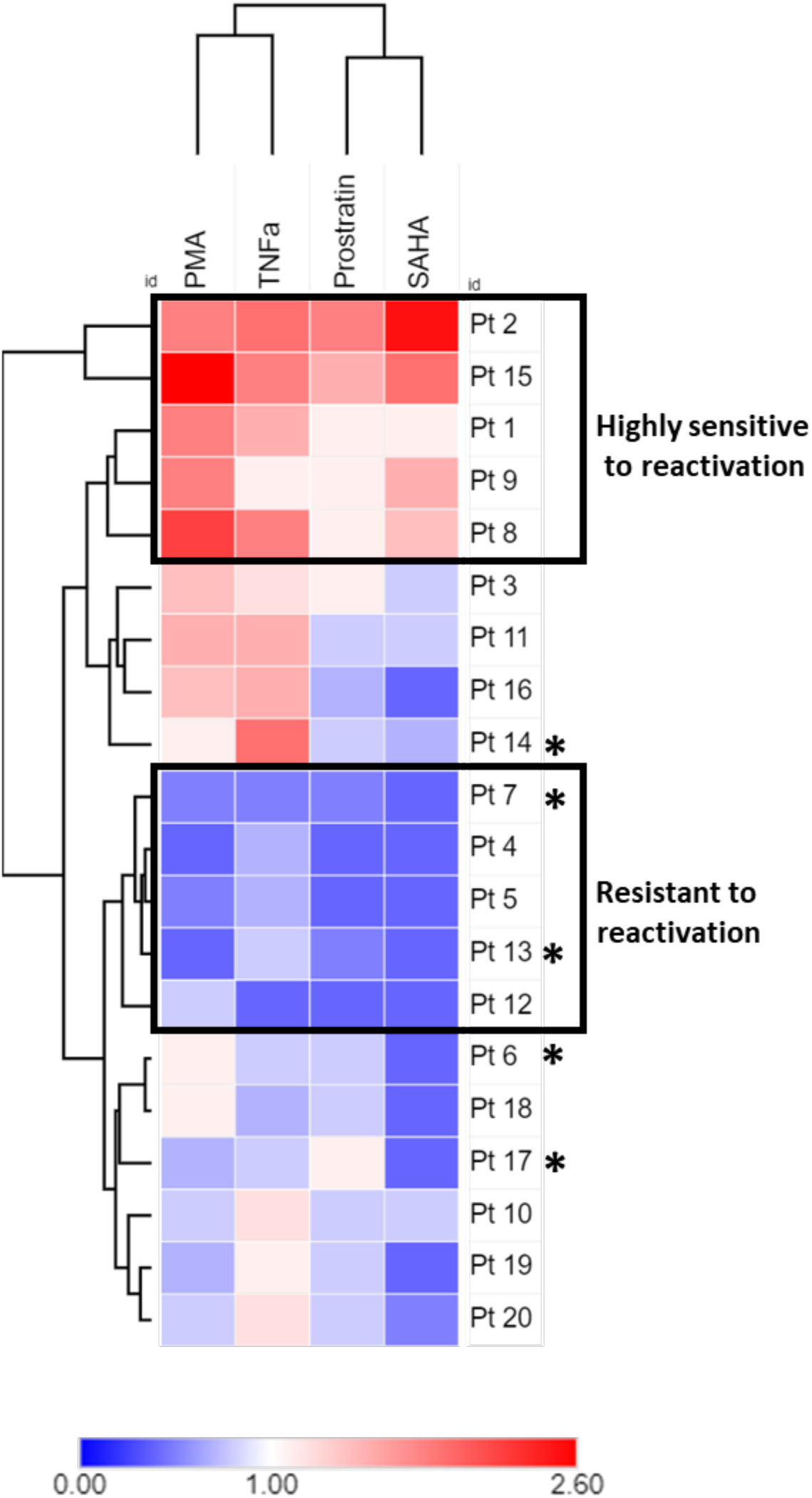
Heatmap showing reactivation of participant LTR-Tat-GFP in latently infected Jurkat cells. Data showing differential reactivation among all 20 participant -derived HIV-1C T/F LTR pseudotyped viruses (denoted as Pt 1-20) when stimulated with LRAs. The viruses that were highly sensitive and resistant to reactivation are depicted within blocks, with the moderately reactivatable viruses remaining outside the blocks. The reactivation percentage of each T/F LTR pseudotyped virus was calculated as the fold change compared to the mean reactivation percentage of all T/F LTR pseudotyped viruses for all treatments. The heatmaps show the dendogram of the hierarchical clustering based on average linkage and the Euclidean distance. The * denotes those participant sequences that contain the 4th NF-kB binding site. The heatmap in this Fig was constructed using the online tool Morpheus (https://software.broadinstitute.org/morpheus).

### Intra-subtype HIV-1 T/F LTR genetic variation impacts reactivation potential

The data from the current study show differential reactivation among PLWH derived HIV-1C T/F LTR pseudotyped latent proviruses. Next, we investigated if differences in latency reversal may be mediated by inter-subtype PLWH-derived HIV-1 T/F LTR genetic variation. For this we grouped the above clusters in 3 groups: highly sensitive, moderately sensitive and resistant to reactivation with LRAs (those highly sensitive and resistant to reactivation are depicted by blocked sections in Fig 3 and coloured sections in Fig 4). Specifically, we observed that latent PLWH derived HIV-1C T/F LTR pseudotyped proviruses that were more sensitive to reactivation by LRAs compared to other variants exhibit only three NF-κB motifs as well as the presence of a Sp1 III cytosine mutation. In contrast, PLWH derived HIV-1C T/F LTR variants that exhibit an insertion of an additional NK-κB (F-kB) (Pt 6,7,13,14 & 17 depicted with an * in Fig 3) just upstream of the AP-1 sequence were moderately or strongly resistant to latency reactivation (Fig 4). Two of those (Pt 7 & 13) are part of the five patient LTRs (40%) that form the strongly resistant cluster of which the remaining three (60%) do not have the fourth NF-κB binding site but are variable within the RBEIII and/or Sp1II sites where the Sp1 III site lacks the cytosine mutation.

**Fig 4:**
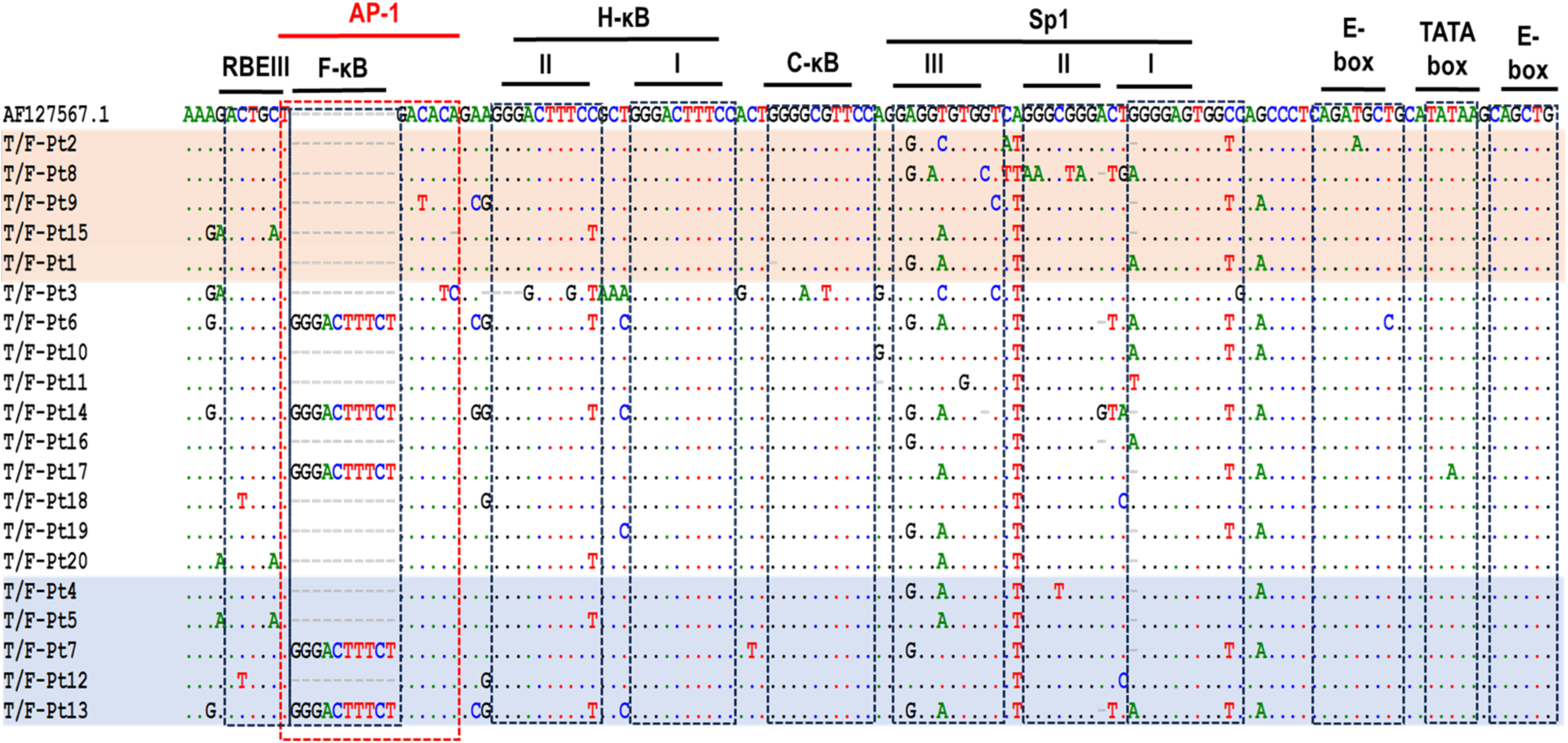
Multiple sequence alignment of participant -derived HIV-1C T/F LTR sequences. The participant derived LTR sequences of highest (highlighted in red) and lowest reactivating variants (highlighted in blue) as determined by unsupervised clustering (Fig 3), were aligned against the African subtype C reference sequence, AF127567.1. The dots within the sequences represent nucleotide bases that are identical to the reference sequence while the grey dashes indicate deletions. The grey dashes within the reference sequence indicate insertions. The dotted blocks highlight the transcription factor binding sites (TFBS) within the core-enhancer and core-promoter regions. From left to right: the AP-1 binding site (enclosed within the red dotted block); the 4th NF-kB binding site (designated F-kB); standard NF-kB sites designated I and II (H-kB, II and I); subtype C specific NF-kB site (designated C-kB); Sp1 III, II and I binding sites; 5’ E-box; Tata box; and 3’ E-box.

Although 3/11 (27%) of the moderately reactivating variants also contain the F-κB, these sequences contain additional polymorphisms that are not present in the variants that were resistant to reactivation, such as T8C mutation within the E box of T/F-Pt6, A8G mutation within the Sp1-II of T/F-Pt14, and T3A mutation within the TATA box of T/F-Pt17 (Fig 4). Consistently with our previous data where we showed that PLWH derived T/F LTR variants impacts transcription activation [20], the data from this study demonstrates that latency reactivation is impacted by LTR genetic variation.

### HIV-1C T/F LTR genetic variation modulates the propensity for latency reversal in primary cells

Following the observation that there is differential reactivation among PLWH derived HIV-1C T/F LTR pseudotyped proviruses in Jurkat E6 cell lines following treatment with different LRAs, we investigated the effect of this genetic variation on the reactivation of latent HIV-1C T/F LTR variants in primary CD4^+^ T-cells. Therefore, we isolated CD4^+^ T-cells from 6 healthy donors (denoted as H 1-6 henceforth) and latently infected these cells with six of the 20 PLWH derived HIV-1C T/F LTR pseudotyped viruses, representing three groups from reactivation in Jurkat E6 cell lines: (1) two highly reactivatable pseudotyped viruses from Pt 2 and Pt 15; (2) two moderately reactivatable pseudotyped viruses from Pt 3 and Pt 11; and (3) two less reactivatable pseudotyped viruses from Pt 4 and Pt 6 (Fig 5 and raw data in Fig S3 and Table S2). Our data showed all 6 PLWH derived HIV-1C T/F LTR pseudotyped latent proviruses exhibited differential reactivation by LRAs. Unsupervised clustering generated roughly eight clusters: one cluster showing high reactivation; two clusters showing low reactivation; and five clusters showing moderate reactivation (Fig 5). Interestingly, we observed that donor variability plays a major role in the reactivation potential of the PLWH derived HIV-1C T/F LTR pseudotyped latent proviruses that differ from the reactivation patterns observed in the Jurkat cells. For example, Pt-4 which is less reactivatable in Jurkat cells is highly reactivatable by PMA and TNF-α in H2 and 3 but not in H4, 5, and 6, while Pt-2 is highly reactivatable in Jurkat cells but is activated poorly by SAHA, Prostratin and TNF-α in H1, 2, 3, and 6 healthy donor cells. This indicates a strong effect of the donor. PMA is able to reactivate 22 out of 36 Patient LTR-Donor combinations, TNF-α 11 out of 36 while prostratin and SAHA reactivate poorly in primary CD4+ healthy donor cells (3 and 1 out of 36 respectively). Interestingly, overall, we find that the LTR derived from Pt6, which is poorly reactivated in J-lats, is most consistently reactivated across conditions and donors (8 out of 24 combinations) and is the only one that is reactivated by all LRAs (in donor H1 which is the donor able to reactivate most Pt LTRs under most conditions (12 out of 24)). This is closely followed by the in J-lats highly activatable Pt2 and 5 LTRs (both 7 out of 24 combinations). Taken together, our data suggest that PLWH HIV-1C T/F LTR genetic variation is a contributing factor in the heterogeneous response to LRAs but also indicate that host cell differences have a major impact on the degree of reactivation induced by a specific LRA.

**Fig 5:**
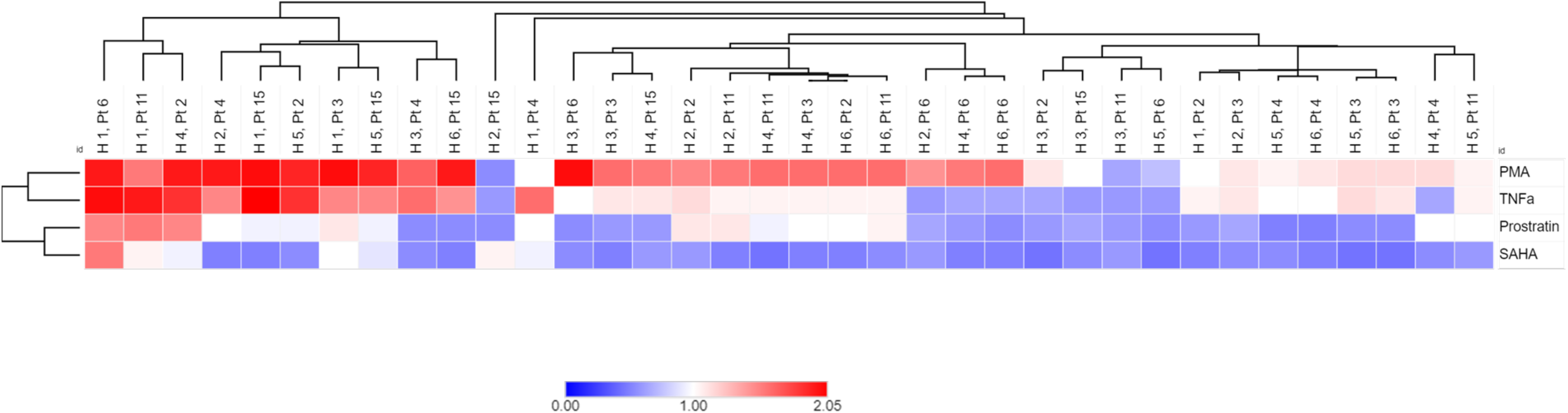
Reactivation of participant LTR-Tat-GFP in latently infected primary cells. All 6 participant -derived HIV-1C T/F LTR pseudotyped viruses showed differential reactivation in all six healthy donor primary cells (denoted as H 1-6) when stimulated with different LRAs. The reactivation percentage of each T/F LTR pseudotyped virus was calculated as the fold change compared to the mean reactivation percentage of all T/F LTR pseudotyped viruses for all treatments. The heatmaps show the dendogram of the hierarchical clustering based on average linkage and the Euclidean distance. The heatmaps in this Fig were constructed using the online tool Morpheus (https://software.broadinstitute.org/morpheus).

### Latent HIV-1C T/F LTR pseudotyped proviruses are sensitive to reactivation by LRAs targeting NF-κB and HDAC pathways albeit at differential levels

Lastly, we hypothesized that the reactivation potentials may differ depending on the LRA used for stimulation, since each LRA works via a different pathway or mechanism to activate the latent virus (reviewed in [4]). To this effect, we performed correlation analysis on the reactivation potentials observed in Jurkat cells. We observed significant positive correlation between reactivation by PMA and TNF-α (r = 0.6469; p < 0.0001) (Fig 6A); PMA and prostratin (r = 0.4452; p = 0.0065) (Fig 6B); TNF-α and prostratin (r = 0.6262; p < 0.0001) (Fig 6D); TNF-α and SAHA (r = 0.4887; p = 0.0025) (Fig 6E); as well as Prostratin and SAHA (r = 0.6136; p<0.0001) (Fig 6F). On the other hand, there was no correlation between reactivation by PMA and SAHA (Fig 6C) suggesting different mechanisms of viral activation by these two LRAs. The HDACi, SAHA, targets the epigenetic environment of the provirus by inhibiting deacetylation of histone tails and transcription repression [9]. Consistent with previous reports that show PKC agonists directly activate viral transcription through NF-ĸB signalling (reviewed in [26]), our data show that PKC agonists PMA, TNF-α and Prostratin used in this study have similar or related mechanisms of viral activation.

**Fig 6:**
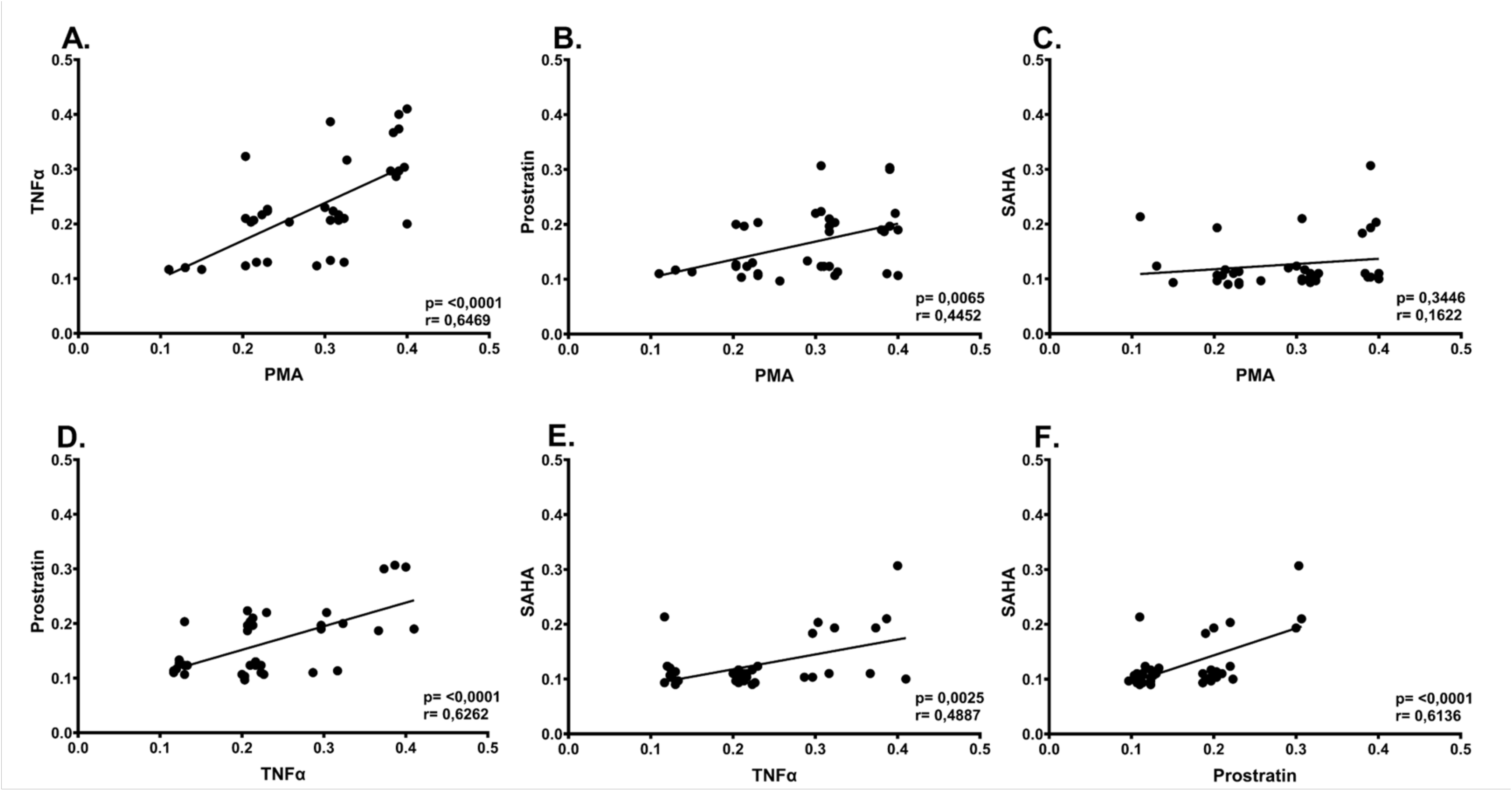
Correlation of reactivation between latency reversing agents. **A:** Correlation plot demonstrating significant positive correlation between PMA and TNF-α (r = 0.6469; p < 0.0001); **B:** Correlation plot demonstrating significant positive correlation between PMA and prostratin (r = 0.4452; p = 0.0065); **C:** Correlation plot demonstrating no correlation between PMA and SAHA; **D:** Correlation plot demonstrating significant positive correlation between TNF-α and prostratin (r = 0.6262; p < 0.0001); **E:** Correlation plot demonstrating significant positive correlation between TNF-α and SAHA (r = 0.4887; p = 0.0025); **F:** Correlation plot demonstrating significant positive correlation between Prostratin and SAHA (r = 0.6136; p<0.0001).

## Discussion

The HIV-1 LTR is the viral promotor that drives viral gene expression, and is important for HIV replication [27]. A previous study reported that genetic variation within the AP-1 motif of the viral promoter regulates the ability of HIV-1 to establish latency [22] while a subsequent study reported that enhanced transcriptional strength of HIV-1C LTR confers stability to the viral latent state [28]. In this study we examined the influence of inter- and intra-subtype LTR genetic variation on the propensity of latency reactivation using different classes of latency reversal agents (LRAs). Our data show that the HIV-1 subtype B exhibited a two-fold higher sensitivity to reactivation with PMA, TNF-α, Prostratin and SAHA compared to subtype C (Fig 1B). Crucially, there were no differences in Tat expression levels or the number of integrated proviral DNA copies between HIV-1B and HIV-1C (Fig 1C & D). However, the role of Tat in HIV-1 latency remains controversial with a previous study reporting that Tat inhibits [29] while the others state that it promotes [23] latency establishment. Additionally, the site of HIV-1 integration in the human genome determines the LTR transcription activity [30]. However, the role of integration sites on latency reactivation was outside of the scope of this study.

A previous study reported that NF-κB p50–HDAC1 complexes constitutively bind the latent HIV-1 LTR and induce histone deacetylation that results in repressive changes in the chromatin structure of the HIV-1 LTR and halts the recruitment of RNA polymerase II and transcriptional initiation [31]. In agreement, our data suggest that subtype B had fewer p50–HDAC1 repressor complexes bound since it had higher reactivation compared to subtype C, which is consistent with the number of NF-κB motifs contained in each subtype. Less reactivation for subtype C in the present study is in line with a report stating an increase in the NF-κB motif copy number stabilizes the latent state [28].

The AP-1 binding site which is upstream of the NF-κB motif within the HIV-1 LTR, controls the establishment of HIV-1 latent infection with its deletion significantly decreasing latency establishment [22]. Interestingly, our findings show that subtype B is associated with enhanced reactivation potential, which could also be attributed to its four-nucleotide AP-1 sequence compared to subtype C seven-nucleotide AP-1 sequence that confers a stable latent viral state.

We previously reported that PLWH derived HIV-1C T/F LTR genetic variation exists and in the current study we show that these PLWH derived HIV-1C T/F LTR pseudotyped proviral variants results in differential reactivation potential in both latently infected Jurkat E6 and primary CD4+ T cells (Fig 3 & 5). Specifically, our findings show three distinct groups of reactivation potential, which were latent proviruses that were highly sensitive, moderately sensitive, and resistant to reactivation by different LRAs (Fig 3 & 4). Interestingly, we noted that latent HIV-1C T/F LTR pseudotyped proviruses that were highly sensitive to reactivation by LRAs all exhibited only three NF-κB sites while the proviruses exhibiting the F-κB site were either moderately reactivatable or resistant to reactivation. This is in line with previous reports showing that a greater number of NF-κB binding sites present within the HIV-1C LTR confers stability to the viral latent state [28]. In most HIV-1C T/F LTRs tested we observed several mutations within the Sp1 III site including A2G, A2G together with T5A, T5C together with T10C, as well as T7G (Fig 4). Previous studies reported that the C-to-T change at nucleotide positions 3 and 5 in the Sp1 III site (designated 3T and 5T respectively) abrogated binding of the Sp1 transcription factor in subtype B LTR [32, 33]. In contrast, a subsequent study reported 3T or 5T mutation alone or 3T5T in combination resulted in moderate or high increase in subtype B LTR-driven gene transcription following TNF-α stimulation [34]. Taken together these data suggest that the specific role of LTR genetic variation within the Sp1 III site remain to be confirmed.

Taken together our data suggests that inter- and intra-subtype LTR genetic variation, in combination with host variation, mediates the propensity of latency reversal by different LRAs.

## Conclusion

We show that inter- and intra-subtype LTR genetic variation mediate the propensity of latency reversal. Specifically, HIV-1 subtype B which contains only two NF-κB motifs was significantly more sensitive to reactivation by different LRAs such as PMA, TNF-α, Prostratin and SAHA compared to subtype C with three NF-κB motifs. Furthermore, our data show that PLWH derived HIV-1C T/F LTR variants exhibiting four NF-κB sites tend, in general, to be less sensitive to reactivation as compared to variants that contained three NF-κB sites. Other mutations in either in RBEIII or Sp1 sites may have also contributed to the heterogenous latency reactivation. Future studies should determine identify cellular biomarkers associated with HIV-1C latency reactivation potential, the role of PLWH derived LTR genetic variation and integration sites on HIV-1C latency establishment or block and lock HIV-1 cure strategy. Our developed PLWH derived transmitted/founder LTR latency model provides an excellent starting point for such studies.

## Materials and Methods

### Ethics statement

The study was approved by the Biomedical Research Ethics Committee (BREC) of the University of KwaZulu-Natal (BREC/00002426/2021). Written informed consent was provided by all study participants.

### Study design and participants

A total of 31 study participants from the Females Rising through Education, Support and Health (FRESH) cohort, and 19 study participants from the HIV Pathogenesis Programme (HPP) Acute Infection cohort were included in this study. FRESH is an ongoing prospective study established in 2012 in Durban, South Africa to study acute HIV-1 infection (AHI) [35, 36]. FRESH recruits 18–23-year-old women who are not living with HIV but are at high risk for infection. FRESH study participants are co-enrolled in 48-week socioeconomic empowerment programme, with classes that coincide with the twice-weekly finger prick blood draw HIV-1 RNA testing. Despite the inclusion of an intensive HIV prevention component and provision of once-daily oral pre-exposure prophylaxis (PrEP), 103 women have become HIV-1 infected since the start of the study. Between April 2013 and June 2014, the first 14 participants were detected during AHI but not started on ART until they met treatment eligibility criteria according to the South African national HIV guidelines (absolute CD4 T-cell count of ≤350 cells/mm^3^). In July 2014, after IRB approval was obtained to provide immediate ART, 89 participants were detected and started immediate ART within a median of 1 day following HIV-1 RNA detection [35]. All participants who acquired HIV-1 infection underwent regular blood and vaginal mucosal sampling. Plasma samples from 9 of the 14 individuals who were not treated during acute infection, as well as from 22 individuals who started ART within 1 to 3 days after HIV-1 RNA detection, were analyzed at the earliest post infection sample timepoint available (pre-ART initiation). The 5’ HIV-1C LTR was amplified and sequenced in these samples, from which 13 were randomly selected to produce the participant -derived HIV-1C T/F LTR pseudotyped viruses (12 individuals who started ART at acute and one individual who started ART at chronic phase) (outlined in Fig S4).

Plasma samples were also obtained from participants in the HPP acute infection cohort in which men and women aged 18–24 years with acute HIV infection were initially enrolled. As we described previously [20], all individuals displayed detectable HIV-1 RNA levels during screening, had not yet seroconverted but subsequently showed an evolving Western blot pattern indicative of recent HIV-1 infection [37]. The date of infection was estimated to be 14 days prior to screening. Blood samples were collected from participants at enrollment, 2 weeks, 4 weeks, 2-, 3- and 6-months post infection and then every 6 months thereafter. The CD4 cell count, and viral loads were determined at each of these visits. The 5’ HIV-1C LTR was amplified and sequenced in 19 plasma samples from the HPP AI cohort obtained at first diagnosis or nearest timepoint depending on sample availability, from which 7 were randomly selected to produce the participant -derived HIV-1C T/F LTR pseudotyped viruses (outlined in Fig 1).

### Construction of subtype C based minimal reporter lentiviral vector “C731CC”

An HIV-1 subtype C based lentiviral vector, C731CC was derived from pEV731 (provided by Dr Tokameh Mahmoudi) where the full-length Tat (both exons one and two of HIV-1B *tat* gene) and GFP are under the control of the HIV-1 subtype B LTR by replacing subtype B LTR and Tat with HIV-1C LTR and Tat respectively (subtype C LTR-Tat-IRES-GFP). Briefly, the HIV-1C consensus LTR (amplified from the following reagent obtained through the NIH HIV Reagent Program, Division of AIDS, NIAID, NIH: Human Immunodeficiency Virus-1 (HIV-1) 93ZM74 LTR Luciferase Reporter Vector, ARP-4789, contributed by Dr. Reink Jeeninga and Dr. Ben Berkhout) and *tat* (amplified from the following reagent obtained through the NIH HIV Reagent Program, Division of AIDS, NIAID, NIH: Plasmid pcDNA3.1(+) Expressing Isogenic Mutant Human Immunodeficiency Virus Type 1 Subtype C BL43/02 Tat (pC-Tat.BL43.CC), ARP-11785, contributed by Dr. Udaykumar Ranga) were purchased as gBlocks (Integrated DNA Technologies) and cloned into the pEV731 HIV minimal reporter genome plasmid devoid of HIV-1B LTR and Tat [30]. The name “C731CC” is derived from subtype C 5’ LTR, 731 (from pEV731), subtype C Tat and subtype C 3’ LTR. Specifically, the gBlock corresponding to the HIV-1C 5’ LTR was digested with *EcoRⅠ*and *BssHⅡ* (New England Biolabs), which were also used to digest the pEV731 vector. The gBlock corresponding to HIV-1C Tat was digested with *ClaⅠ* and *BamHⅠ* (New England Biolabs) which were also used to digest pEV731. The gBlock corresponding to HIV-1C 3’ LTR was digested with *PacⅠ* and *XhoⅠ* (New England Biolabs) and these enzymes were also used to restrict the pEV731 vector. The restriction fragments of the pEV731 plasmid were analyzed on a 1% agarose gel and the bigger band fragment containing the linearized plasmid was extracted using the GeneJet Gel Extraction Kit (ThermoFisher Scientific) as per manufacturer’s instructions. The corresponding LTR and Tat gBlocks were cloned into the linearized pEV731 vector lacking either 5’ LTR, Tat or 3’ LTR by ligation using 1 U of T4 DNA ligase (New England Biolabs) as per manufacturer’s instructions, to generate the recombinant lentiviral minimal reporter construct for subtype C, “C731CC”. NEB® Stable Competent *E. coli* cells (High Efficiency) (New England Biolabs) were then transformed with this C731CC vector as per manufacturer’s instructions and grown overnight for 14 hours at 30°C on ampicillin agar plates. The C731CC vector (plasmid map shown in Fig S5) was then purified from the bacterial cells using the GeneJet Plasmid Mini Prep Kit (Invitrogen) as per the manufacturer’s instructions.

### HIV-1C participant -derived transmitted/founder (T/F) LTR amplification, sequencing, and relatedness analysis

A total of 50 participant -derived HIV-1C transmitted/founder (T/F) U3R regions of the LTR contained in the recombinant pGL3 Basic vector [20] (referred to as T/F LTR hence forth) were amplified using forward primer (LTRC_*ECORI*-F: 5’- TAA TAC GAC TCA CTA TAG GGT T<u>GA ATT C</u>TT TAA AAG AAA AGG GGG GAC -3’) containing the *EcoRI* site (underlined) and reverse primer (LTRC_*BssHII*-R: 5’- ATT TAG GTG ACA CTA TAG AGC TTT ATT GAG GC<u>G CGC GC</u>A GTG GGT T -3’) containing the *BssHII* site (underlined). The polymerase chain reaction (PCR) was performed using the Platinum™ Taq DNA Polymerase High Fidelity PCR Kit (Invitrogen), according to the manufacturer’s instructions. Briefly, a PCR reaction was prepared on ice with 1X High Fidelity Buffer; 2 mM MgSO_4_; 0.2 mM dNTP Mix; 0.2 μM forward primer; 0.2 μM reverse primer; 1 ng template DNA; 1U of Platinum® Taq DNA Polymerase High Fidelity enzyme and PCR-grade water to make it up to a final reaction volume of 25 µL. PCR cycling conditions included an initial denaturation for 5 minutes at 94°C, followed by 35 cycles of denaturation at 94°C for 15 seconds; annealing at 55°C for 30 seconds; extension at 68°C for 30 seconds, followed by a final extension of 68°C for 7 minutes. PCR products were then analyzed on 1% agarose gel and purified using the QIAquick PCR Purification Kit (Qiagen, Valencia, CA) according to the manufacturer’s instructions.

This purified PCR product and the corresponding HIV-1C T/F LTR still contained in the recombinant pGL3 Basic vector were sequenced using the BigDye™ Terminator v3.1 Cycle Sequencing Kit (ThermoFisher Scientific). Briefly, a sequencing reaction was separately prepared for each primer containing 2 µL of 0.4 μM primer (forward or reverse primer); 2μL sequencing buffer; 1 μL of 20 ng/μL DNA template; 0.4 μL of BigDye v3.; and 3.4 μL of PCR grade water. This reaction was then centrifuged and subjected to an initial denaturation at 96°C for 1 minute, 25 cycles of denaturation at 96°C for 10 seconds, annealing at 50°C for 5 seconds, and extension at 50°C for 4 seconds. This was followed by a sequencing reaction clean up, in which 1 µL of 125 mM ethylenediaminetetraacetic acid (EDTA) (pH 8.0); 1 µL of 3 M sodium acetate (NaOAc) (pH 5.2); and 25 µL of 100% ethanol were added to each reaction, vortexed and centrifuged for 20 minutes at 3,000 rpm. The supernatant was then removed by inverting the reaction and centrifuging again for 1 minute. This was followed by the addition of 35 μL of 70% ethanol to each reaction, centrifugation for 5 minutes (3,000 rpm), and removal of the supernatant again by inverting the reaction and centrifuging for another minute. The sequencing reactions were then dried at 50°C for 5 minutes and stored at 4°C with protection from exposure to light. These samples were then analyzed using the ABI 3130xl Genetic Analyzer. All the HIV-1 LTR sequences were assembled and analysed using the Sequencher Program v5.0 (Gene Codes Corporation). Phylogenetic relatedness analysis to compare and evaluate the similarity between pGL3 Basic vector and C731CC lentiviral vector derived sequences was performed by Neighbor-Joining trees (with 1,000 bootstrap replicates) using PhyML Maximum Likelihood software (https://www.hiv.lanl.gov/content/sequence/PHYML/interface.html). Branching topology was visualized in Figtree (http://tree.bio.ed.ac.uk/software/figtree). Multiple sequence alignment was done using MAFFT and visualized using BioEdit. The South African HIV-1C reference strain was obtained from the Los Alamos HIV sequence database (www.hiv.lanl.gov).

### Generation of participant -derived T/F LTR -C731CC minimal reporter constructs

The participant derived HIV-1 T/F LTR was then cloned into the C731CC vector to create a C_T/F_731CC. Briefly, the purified HIV-1C T/F LTR PCR products and C731CC were digested with *EcoRI* and *BssHII* (New England Biolabs). The restriction fragments of the C731CC lentiviral vector were analyzed on the 1% agarose gel and the bigger band fragment containing the linearized C731CC lentiviral vector was extracted using the GeneJet Gel Extraction Kit (ThermoFisher Scientific) as per manufacturer’s instructions. The purified HIV-1C T/F LTR PCR products were cloned into the linearized C731CC lentiviral vector by ligation using 1 U of T4 DNA ligase (New England Biolabs) as per manufacturer’s instructions, to generate C_T/F_731CC minimal reporter lentiviral vector. NEB® Stable Competent *E. coli* cells (High Efficiency) (New England Biolabs) were then transformed with C_T/F_731CC minimal genome reporter lentiviral vector as described above. The C_T/F_731CC minimal genome reporter lentiviral vectors were then purified from the bacterial cells using the GeneJet Plasmid Mini Prep Kit (Invitrogen) as described above. An overview of this methodology is shown in Fig 2B.

### Cell culture

Human embryonic kidney (HEK) 293T cells were cultured in Dulbecco’s Modified Eagle’s Medium (DMEM) (ThermoFisher Scientific) supplemented with 10% fetal bovine serum (FBS), 100 μg/ml penicillin-streptomycin, and 1% HEPES at 37°C in a humidified 95% air- 5% CO_2_ atmosphere. Jurkat cells were grown in RPMI 1640 medium containing L-glutamine (ThermoFisher Scientific) supplemented with 10% FBS, 100 μg/ml penicillin-streptomycin, and 1% HEPES (to make R10 medium) at 37°C in a humidified 95% air-5% CO_2_ atmosphere.

### Production of HIV-1C LTR pseudotyped viruses

The HEK 293T cells were co-transfected with the C731CC or C_T/F_731CC minimal reporter lentiviral vector (plasmid); the vesicular stomatitis virus (VSV-G) plasmid to contribute the pseudotyped Envelope, as well as the pCMVΔR8.91 packaging vector which provides all required vector proteins. Briefly, on day zero 2×10^6^ HEK 293T cells were seeded (in supplemented DMEM as described above) in a T75 flask to achieve ∼70% confluency by the next day. On day one, the medium was changed to DMEM only (no supplements) and a DNA mix comprising of 6 µg C731CC vector, 2 µg VSV-G plasmid, and 4.5 µg pCMVΔR8.91 packaging vector in a total of 500 µL was made. A transfection mix was made with 125 µL Polyethylenimine (PEI) Transfection Reagent (ThermoFisher Scientific) and DMEM to a total of 500 µL followed by incubation at room temperature for 5 minutes. The transfection mix was then added to the DNA mix dropwise and incubated at room temperature for 20 minutes. This PEI/DNA mix was then added dropwise to the cells in the T75 flask and incubated for 6 hours at 37°C in a humidified 95% air-5% CO_2_ atmosphere followed by a media change to R10 medium. The pseudotyped viruses were then harvested from the supernatant of cell cultures at 72 hours post media change, filter sterilized with a 0.45 μm filter, aliquoted, and stored at −80°C. An overview of this methodology is shown in Fig 2B.

### Jurkat cell infection

The Jurkat cells were grown to have at least 95% viability and were infected with C731CC or C_T/F_731CC pseudotyped viruses, such that approximately 5% of cells were infected. Briefly, 4 x 10^5^ cells/mL were seeded in a 6 well plate and infected with C731CC or C_T/F_731CC pseudotyped virus for 96 hours of incubation at 37°C in a humidified 95% air-5% CO_2_ atmosphere to give about 5% GFP positive cells. An overview of this methodology is shown in Fig 2B.

### Flow cytometry

GFP expression was analyzed by flow cytometry. The live population was defined by forward versus side scatter profiles. Gating for SSC-H vs SSC-W and FSC-H vs FSC-W was used to exclude doublets. Cells were further gated by using forward scatter versus GFP intensity to differentiate between GFP-positive and -negative cells (gating strategy shown in Fig S6).

The 95% of GFP negative cell population, which represented either true negative or latently infected cells were sorted using a BD FACSAria™ Fusion Flow Cytometer (Becton Dickinson) and treated with latency reversing agents (LRAs).

### Latency reversal

Immediately following cell sorting of the GFP negative cells, the cells were centrifuged at 1,500 rpm for 10 minutes and resuspended in R10 medium. The following LRAs, 2 mM phorbol 12-myristate 13-acetate (PMA); 2 µg/mL tumor necrosis factor alpha (TNF-α); 2 µg/mL prostratin; and 1500 nM suberoylanilide hydroxamic acid (SAHA) were added. The concentrations of the LRAs used are a representative concentration that was not toxic to the cell viability as viability remained high above 90%. This was followed by 24 hours incubation at 37°C in a humidified 95% air-5% CO_2_ atmosphere. The percentage reactivation was then measured as the percentage GFP positive cells by flow cytometry as described above. These experiments were performed in triplicates.

### Western blot

J-Lat and C J-Lat cells were lysed with IP buffer (1% NP40, 5% glycerol, 5 mM MgCl_2_, 1 mM EDTA, 150 mM KCl, 25 mM HEPES, pH 7.9, 0.5 mM dithiothreitol and a protease inhibitor cocktail) (Sigma Aldrich) on ice for 30 minutes. Whole-cell protein lysate was used for SDS-PAGE to detect Flag-tagged HIV-1 subtype B Tat (Tat B) and HIV-1 subtype C Tat (Tat C) with anti-Flag antibody (Sigma Aldrich) and anti-mouse IgG polyclonal HRP antibody (Bio-Rad). Tubulin was used as a loading control.

### Measurement of the amount of integrated HIV-1 DNA copies

Quantification of the integrated HIV-1 DNA copies was performed as previously described [38]. DNA was isolated from both J-Lat and C J-Lat cells by lysing 2 x 10^6^ cells using the AllPrep DNA/RNA Mini Kit (Qiagen) as per the manufacturer’s instructions. Alu-gag PCR was then conducted as follows: The first round PCR reaction was prepared on ice using 1X PCR buffer (ThermoFisher), 1.5 mM MgSO_4_ (ThermoFisher), 0.5 mM dNTPs (ThermoFisher), 1 μM Alu forward primer (5’-GCC TCC CAA AGT GCT GGG ATT ACA G-3’), 6 μM Gag reverse primer (5’-TCG CTT TCA GGT CCC TGT TCG-3’), 2.5 U of Platinum Taq polymerase (ThermoFisher), and 150 ng of genomic DNA. Cycling conditions were as follows: initial denaturation of 95°C for 2 minutes, followed by 14 cycles of denaturation at 95°C for 30 seconds; annealing at 50°C for 30 seconds and extension at 72°C for 210 seconds. The nested PCR reaction was then prepared on ice using 1X PCR buffer (ThermoFisher), 1.5 mM MgSO_4_ (ThermoFisher), 0.5 mM dNTPs (ThermoFisher), 0.5 μM AluGag forward primer (5’-GGT GCG AGA GCG TCA GTA T-3’) and 0.5 μM AluGag reverse primer (5’-AGC TCC CTG CTT GCC CAT A-3’), 0.15 μM AluGag probe (6FAM-AAA ATT CGG TTA AGG CCA GGG GGA AAG AA-QSY7), 1 U of Platinum Taq polymerase (ThermoFisher), and 2 μL of Alu-gag PCR product. Cycling conditions were as follows: 95 °C for 5 minutes, followed by 50 cycles of 95°C for 10 seconds and 60 °C for 30 seconds. Human ribonuclease (RNase) P was used for normalization (ThermoFisher).

### Primary CD4^+^ T-Cell Isolation and Infection

Peripheral blood mononuclear cells (PBMCs) were isolated from healthy donors by Ficoll gradient followed by primary CD4^+^ T-cell isolation by negative selection and magnetic separation using the human CD4^+^ T Cell Isolation kit (Miltenyi Biotec) according to the manufacturer’s instructions. Isolated CD4^+^ T-cells were then rested in R10 medium at a concentration of 1×10^6^ cells/mL for 3 hours at 37°C in a humidified 95% air-5% CO_2_ atmosphere before HIV-1 infection. The CD4^+^ T-cells were then centrifuged at 800 x g and resuspended in 200 µL of viral supernatant in a 15 mL conical centrifuge tube. The spinoculation was then carried out for 2 hours at 1200 x g at room temperature. After spinoculation, the cells were cultured in R10 medium supplemented with 5 mM Saquinavir to prevent residual spreading infection for 72 hours at 37°C in a humidified 95% air-5% CO_2_ atmosphere. Saquinavir was obtained through the AIDS Research and Reference Reagent Program, Division of AIDS, NIAID, NIH. The cells were then treated with LRAs in presence of 30 mM Raltegravir (obtained from the AIDS Research and Reference Reagent Program, Division of AIDS, NIAID, NIH) for 24 hours, followed by measurement of GFP positive cells by flow cytometry as described above, to determine the percentage reactivation. These experiments were performed in triplicates. The 2μM concentration of the PMA used is a representative concentration that was not toxic to the cells as viability remained high above 99%.

### Statistical and heatmap analysis

Statistical analysis was performed using GraphPad Prism 10 software. The statistical significances for the B731BB and C731CC latency reactivation comparisons were determined using an unpaired T-test, while that for integrated DNA copy measurements were determined using a non-parametric test. Linear regression analysis was used to determine the correlation coefficient and statistical significance of the correlation between the different LRAs. A p-value ≤ 0.05 was considered statistically significant. The heatmaps were constructed using the online tool Morpheus (https://software.broadinstitute.org/morpheus). The reactivation percentage of each T/F LTR pseudotyped virus was calculated as the fold change compared to the mean reactivation percentage of all T/F LTR pseudotyped viruses for all treatments. The heatmaps show the dendogram of the hierarchical clustering based on average linkage and the Euclidean distance.

## Acknowledgments

The authors would like to acknowledge the study participants and the clinical teams from the FRESH and HIV Pathogenesis Programme (HPP) acute infection studies.

Research reported in this publication was supported by the South African Medical Research Council with funds received from the National Department of Health (MRC-RFA-SHIP 02-2018 to PM) and a grant from the National Research Foundation Thuthuka Funding Instrument (TTK160529166617 to PM and PMDS22070735179 to SM). This study was funded by the Poliomyelitis Research Foundation (Grant number: 21/57). The work was also supported in part by grants from the Bill and Melinda Gates Foundation [OPP1212883 and INV-033558]; Gilead Sciences, Inc [Grant ID #00406] and the International AIDS Vaccine Initiative (IAVI) [UKZNRSA1001 to TN]. This work also received funding through the Sub-Saharan African Network for TB/HIV Research Excellence (SANTHE) [grant # DEL-15-006]. SANTHE receives additional funds from the Science for Africa Foundation with support from Wellcome Trust and the UK Foreign, Commonwealth & Development Office and is part of the EDCPT2 programme supported by the European Union; the Bill & Melinda Gates Foundation [INV-033558]; and Gilead Sciences Inc., [19275]. All content contained within is that of the authors and does not necessarily reflect positions or policies of any SANTHE funder. For the purpose of open access, the author has applied a CC BY public copyright licence to any Author Accepted Manuscript version arising from this submission. Lastly, the study was also partly funded by Erasmus+ which is the EU’s programme to support education, training, youth and sport in Europe [Erasmus+KA107].

## Supporting information captions

**Table S1:** Raw reactivation values of patient LTR-Tat-GFP in latently infected Jurkat cells as shown in heatmap in Fig 3.

**Table S2:** Raw reactivation values of patient LTR-Tat-GFP in latently infected primary cells as shown in heatmap in Fig 5.

**Fig S1: Sequence alignment of consensus HIV-1 subtype B and C LTR core enhancer.** The HIV-1 subtype B consensus 5’ LTR contains a four-nucleotide AP-1 motif just upstream of the NF-κB element, while the HIV-1 subtype C consensus 5’ LTR contains an extension of this AP-1 motif to seven nucleotides (highlighted in yellow).

**Fig S2: Bar graph depicting reactivation of patient LTR-Tat-GFP in latently infected Jurkat cells as shown in heatmap in Fig 3**. All 20 patient-derived HIV-1C T/F LTR pseudotyped viruses (denoted as Pt 1-20) reactivated significantly following addition of the LRAs alone or in combination (denoted by +) compared to the unstimulated control (black bars denoted by -). Furthermore, there was differential reactivation among the patient-derived HIV-1C T/F LTR pseudotyped viruses when stimulated with different LRAs. The two highly reactivating viruses from Pt 2 and Pt 15 are denoted by the turquoise upward arrows; two moderately reactivating viruses from Pt 3 and Pt 11 are denoted by the maroon sideward arrows; and two low reactivating viruses from Pt 4 and Pt 6 are denoted by the orange downward arrows. **A**: Reactivation potentials of all 20 viruses upon PMA stimulation. **B**: Reactivation potentials of all 20 viruses upon TNF-α stimulation. **C**: Reactivation potentials of all 20 viruses upon prostratin stimulation. **D**: Reactivation potentials of all 20 viruses upon SAHA stimulation.

**Fig S3: Bar graph depicting reactivation of patient LTR-Tat-GFP in latently infected primary cells as shown in heatmap in Fig 5**. All 6 patient-derived HIV-1C T/F LTR pseudotyped viruses reactivated significantly above the threshold dotted line following addition of the latency reversing agents compared to the unstimulated control (black bars) in all six healthy donor primary cells. PMA stimulation is denoted by red bars, TNF-α stimulation is denoted by blue bars, prostratin stimulation is denoted by green bars, and SAHA stimulation is denoted by purple bars. There was differential reactivation among the patient-derived HIV-1C T/F LTR pseudotyped viruses when stimulated with different LRAs in all six healthy donor primary cells.

**Fig S4: South African acute HIV-1 infection cohort design and the number of participants from each cohort used to generate HIV-1C T/F LTR sequences**. Initiated ART at acute refers to the study participants that initiated treatment shortly after viral RNA detection. Initiated ART at chronic refers to the study participants who initiated treatment when their CD4 T cell count was below 500 cells/μL according to South African national treatment guidelines at the time.

**Fig S5: Plasmid map of the recombinant lentiviral minimal reporter construct for subtype C, “C731CC”.** Adapted from the pEV731 HIV minimal reporter genome plasmid by cloning a HIV-1C consensus LTR and tat in place of the HIV-1B LTR (both 5’ and 3’) and tat in the pEV731 lentiviral vector, respectively. In the C731CC plasmid (subtype C LTR-Tat-IRES-EGFP), the expression of HIV-1C Tat and GFP are under the control of the HIV-1C 5’ LTR.

**Fig S6: Gating strategy used for flow cytometry.** The live population was defined by forward versus side scatter profiles (P1). Gating for SSC-H vs SSC-W (P2) and FSC-H vs FSC-W (P3) was used to exclude doublets. Cells were further gated by using forward scatter versus GFP intensity to differentiate between GFP-negative (P4) and -positive cells (P5).

