## Supplemental Data for "Development of a latency model based on HIV-1 subtype C to study how long terminal repeat genetic variation impacts viral persistence and latency reversal"

**Supporting information:**

| | Unstimulated | PMA | TNF-α | Prostratin | SAHA |
|---|---|---|---|---|---|
| **Pt 1** | 0.00 | 0.48 | 0.40 | 0.31 | 0.29 |
| **Pt 2** | 0.01 | 0.49 | 0.51 | 0.49 | 0.68 |
| **Pt 3** | 0.01 | 0.39 | 0.32 | 0.30 | 0.21 |
| **Pt 4** | 0.01 | 0.11 | 0.20 | 0.12 | 0.11 |
| **Pt 5** | 0.01 | 0.13 | 0.19 | 0.12 | 0.11 |
| **Pt 6** | 0.01 | 0.29 | 0.22 | 0.21 | 0.11 |
| **Pt 7** | 0.01 | 0.12 | 0.13 | 0.13 | 0.11 |
| **Pt 8** | 0.01 | 0.59 | 0.49 | 0.31 | 0.39 |
| **Pt 9** | 0.01 | 0.49 | 0.30 | 0.29 | 0.40 |
| **Pt 10** | 0.01 | 0.21 | 0.32 | 0.23 | 0.20 |
| **Pt 11** | 0.01 | 0.42 | 0.40 | 0.20 | 0.21 |
| **Pt 12** | 0.02 | 0.21 | 0.12 | 0.11 | 0.11 |
| **Pt 13** | 0.01 | 0.12 | 0.20 | 0.13 | 0.12 |
| **Pt 14** | 0.01 | 0.31 | 0.52 | 0.20 | 0.19 |
| **Pt 15** | 0.01 | 0.71 | 0.49 | 0.41 | 0.50 |
| **Pt 16** | 0.02 | 0.38 | 0.42 | 0.18 | 0.12 |
| **Pt 17** | 0.01 | 0.19 | 0.21 | 0.30 | 0.11 |
| **Pt 18** | 0.01 | 0.29 | 0.20 | 0.21 | 0.10 |
| **Pt 19** | 0.01 | 0.19 | 0.31 | 0.20 | 0.12 |
| **Pt 20** | 0.00 | 0.21 | 0.32 | 0.22 | 0.13 |

### Table S1

| Unstimulated | | | | | | |
|---|---|---|---|---|---|---|
| | **H1** | **H2** | **H3** | **H4** | **H5** | **H6** |
| **Pt 2** | 0.02 | 0.02 | 0.02 | 0.00 | 0.02 | 0.02 |
| **Pt 3** | 0.01 | 0.00 | 0.01 | 0.01 | 0.01 | 0.01 |
| **Pt 4** | 0.01 | 0.02 | 0.03 | 0.02 | 0.01 | 0.01 |
| **Pt 6** | 0.01 | 0.01 | 0.02 | 0.01 | 0.02 | 0.02 |
| **Pt 11** | 0.01 | 0.01 | 0.01 | 0.01 | 0.02 | 0.02 |
| **Pt 15** | 0.03 | 0.02 | 0.01 | 0.01 | 0.02 | 0.02 |
| **PMA** | | | | | | |
| | **H1** | **H2** | **H3** | **H4** | **H5** | **H6** |
| **Pt 2** | 0.20 | 0.30 | 0.22 | 0.39 | 0.38 | 0.32 |
| **Pt 3** | 0.40 | 0.22 | 0.32 | 0.32 | 0.23 | 0.23 |
| **Pt 4** | 0.20 | 0.39 | 0.33 | 0.23 | 0.21 | 0.22 |
| **Pt 6** | 0.39 | 0.29 | 0.40 | 0.31 | 0.15 | 0.32 |
| **Pt 11** | 0.31 | 0.31 | 0.13 | 0.32 | 0.21 | 0.32 |
| **Pt 15** | 0.40 | 0.11 | 0.20 | 0.31 | 0.38 | 0.39 |
| **TNF-α** | | | | | | |
| | **H1** | **H2** | **H3** | **H4** | **H5** | **H6** |
| **Pt 2** | 0.21 | 0.23 | 0.13 | 0.37 | 0.37 | 0.21 |
| **Pt 3** | 0.30 | 0.22 | 0.22 | 0.21 | 0.23 | 0.22 |
| **Pt 4** | 0.32 | 0.30 | 0.32 | 0.13 | 0.20 | 0.20 |
| **Pt 6** | 0.40 | 0.12 | 0.20 | 0.13 | 0.12 | 0.13 |
| **Pt 11** | 0.39 | 0.21 | 0.12 | 0.21 | 0.21 | 0.21 |
| **Pt 15** | 0.41 | 0.12 | 0.12 | 0.22 | 0.30 | 0.29 |
| **Prostratin** | | | | | | |
| | **H1** | **H2** | **H3** | **H4** | **H5** | **H6** |
| **Pt 2** | 0.12 | 0.22 | 0.12 | 0.30 | 0.19 | 0.20 |
| **Pt 3** | 0.22 | 0.13 | 0.12 | 0.20 | 0.11 | 0.11 |
| **Pt 4** | 0.20 | 0.20 | 0.11 | 0.20 | 0.10 | 0.10 |
| **Pt 6** | 0.30 | 0.13 | 0.11 | 0.12 | 0.11 | 0.11 |
| **Pt 11** | 0.31 | 0.22 | 0.12 | 0.19 | 0.20 | 0.21 |
| **Pt 15** | 0.19 | 0.11 | 0.13 | 0.12 | 0.19 | 0.11 |
| **SAHA** | | | | | | |
| | **H1** | **H2** | **H3** | **H4** | **H5** | **H6** |
| **Pt 2** | 0.10 | 0.12 | 0.09 | 0.19 | 0.11 | 0.10 |
| **Pt 3** | 0.20 | 0.11 | 0.10 | 0.10 | 0.09 | 0.09 |
| **Pt 4** | 0.19 | 0.10 | 0.11 | 0.11 | 0.11 | 0.10 |
| **Pt 6** | 0.31 | 0.12 | 0.11 | 0.10 | 0.09 | 0.10 |
| **Pt 11** | 0.21 | 0.10 | 0.12 | 0.09 | 0.12 | 0.11 |
| **Pt 15** | 0.10 | 0.21 | 0.11 | 0.12 | 0.18 | 0.10 |

### Table S2

**AP-1**

**Subtype B consensus**: TTCAAGAACTGC**TGAC**ATCGAGCT TGCT .ACA .. AGGGACTTTCCGCTGGGGACTTTCC
**Subtype C consensus**: GGGACTTTCTGC**TGACACA**GAAGGGACTTTCCGCTGGGACTTTCCACCG .GGGCGTTCC

### Figure S1

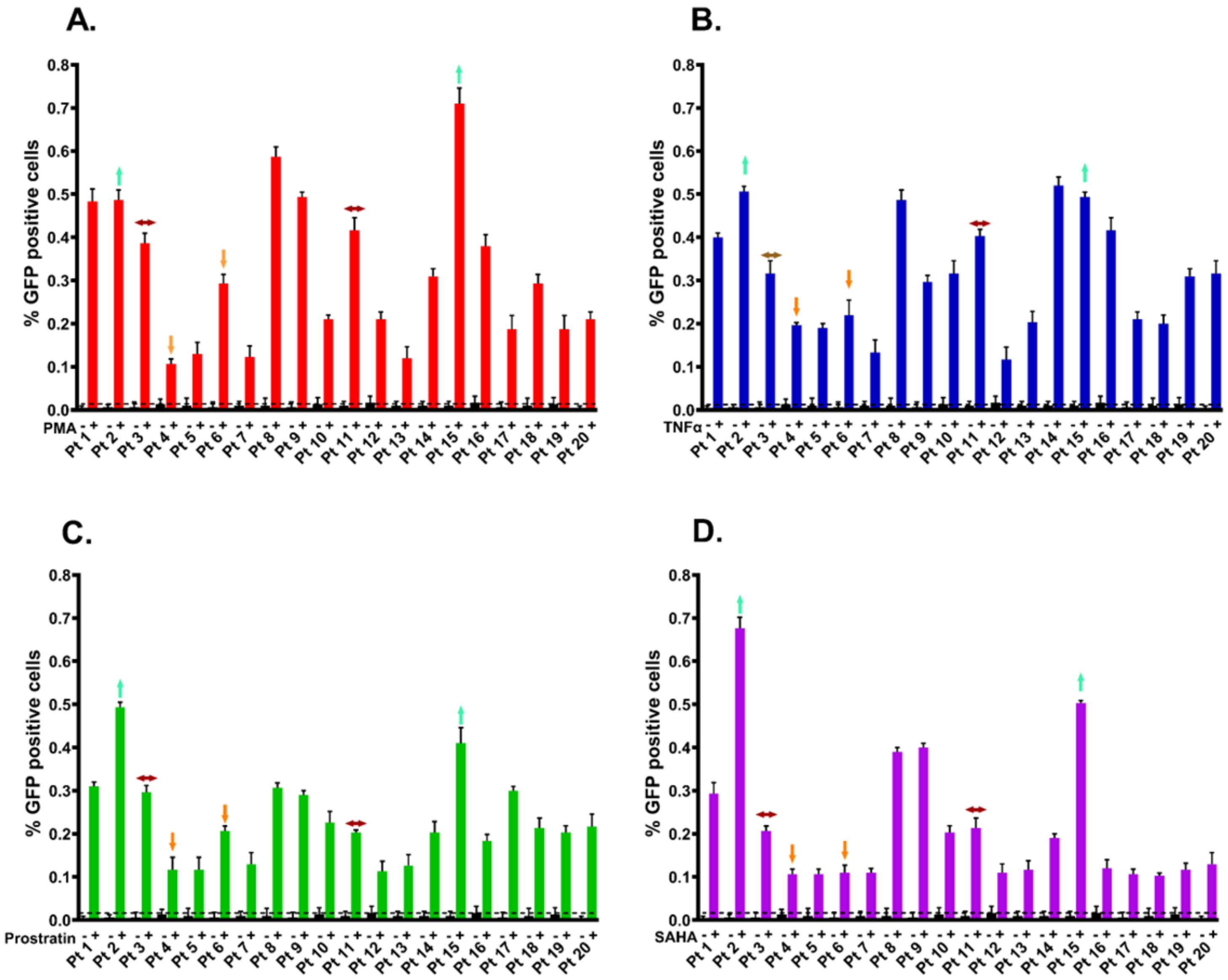
A.
B.
C.
D.
% GFP positive cells
PMA
TNFα
Prostratin
SAHA
Pt 1
Pt 2
Pt 3
Pt 4
Pt 5
Pt 6
Pt 7
Pt 8
Pt 9
Pt 10
Pt 11
Pt 12
Pt 13
Pt 14
Pt 15
Pt 16
Pt 17
Pt 18
Pt 19
Pt 20
0.0
0.1
0.2
0.3
0.4
0.5
0.6
0.7
0.8

### Figure S2

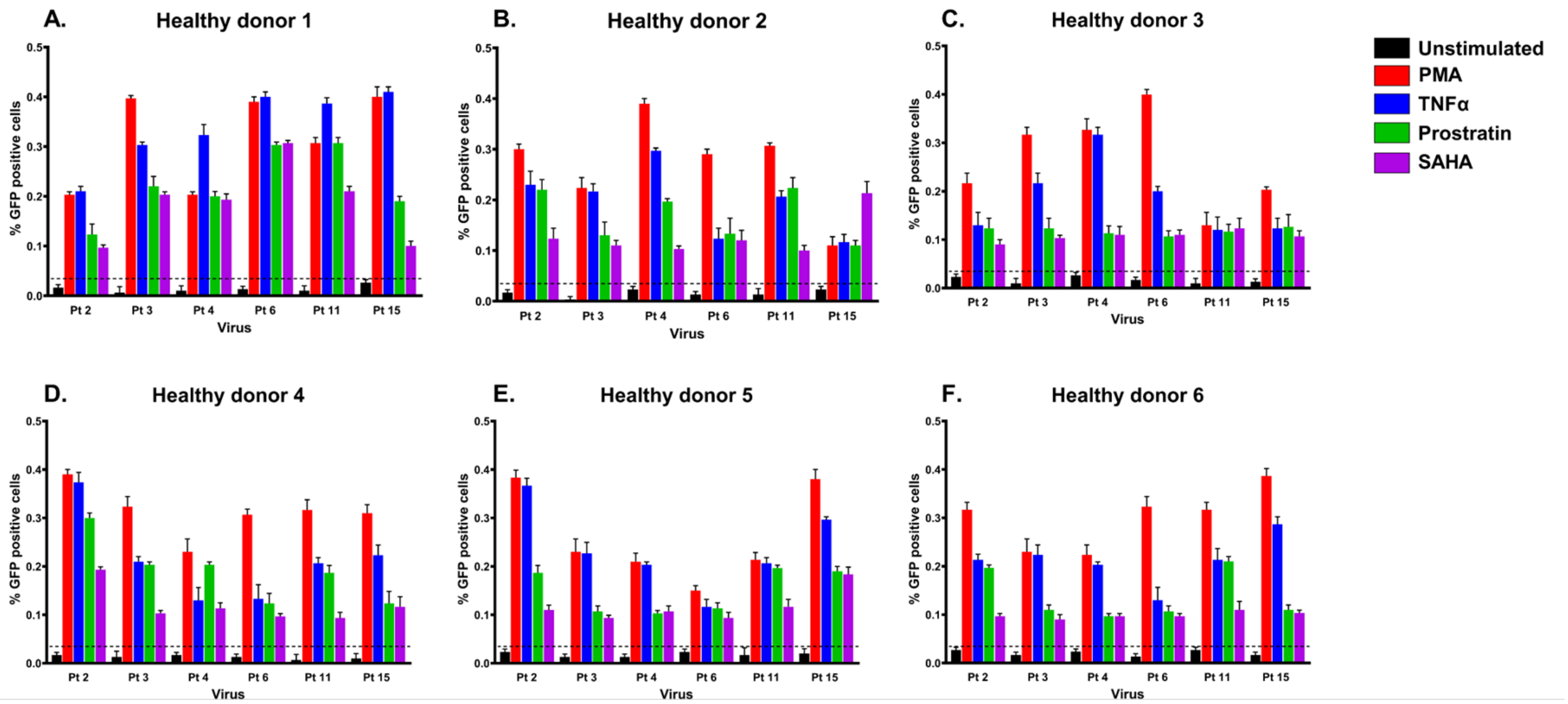
A.
Healthy donor 1
B.
Healthy donor 2
C.
Healthy donor 3
D.
Healthy donor 4
E.
Healthy donor 5
F.
Healthy donor 6
% GFP positive cells
Virus
Pt 2
Pt 3
Pt 4
Pt 6
Pt 11
Pt 15
Unstimulated
PMA
TNFα
Prostratin
SAHA

### Figure S3

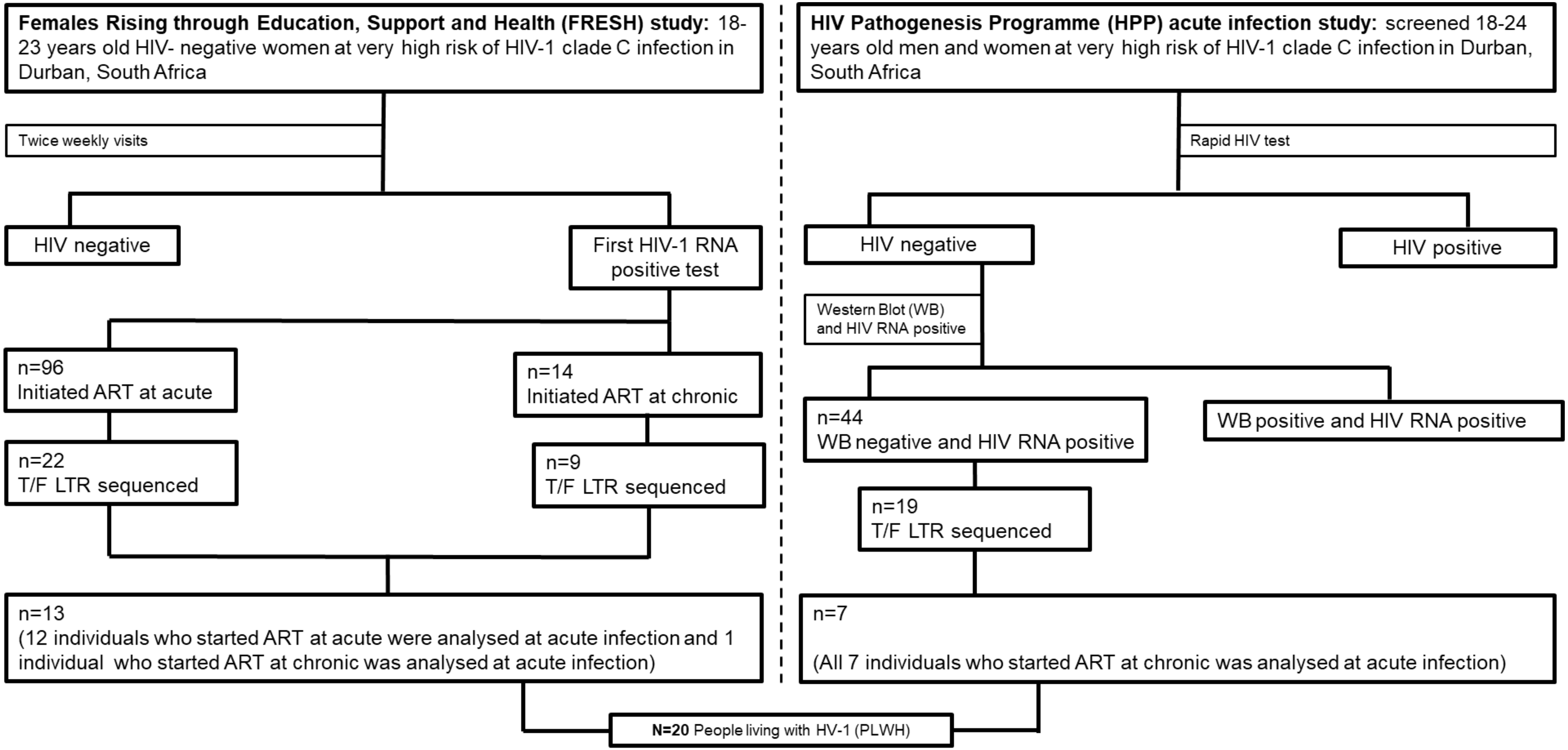
Females Rising through Education, Support and Health (FRESH) study: 18-23 years old HIV- negative women at very high risk of HIV-1 clade C infection in Durban, South Africa
Twice weekly visits
HIV negative
First HIV-1 RNA positive test
n=96 Initiated ART at acute
n=14 Initiated ART at chronic
n=22 T/F LTR sequenced
n=9 T/F LTR sequenced
n=13 (12 individuals who started ART at acute were analysed at acute infection and 1 individual who started ART at chronic was analysed at acute infection)
HIV Pathogenesis Programme (HPP) acute infection study: screened 18-24 years old men and women at very high risk of HIV-1 clade C infection in Durban, South Africa
Rapid HIV test
HIV negative
HIV positive
Western Blot (WB) and HIV RNA positive
n=44 WB negative and HIV RNA positive
WB positive and HIV RNA positive
n=19 T/F LTR sequenced
n=7
(All 7 individuals who started ART at chronic was analysed at acute infection)
N=20 People living with HV-1 (PLWH)

### Figure S4

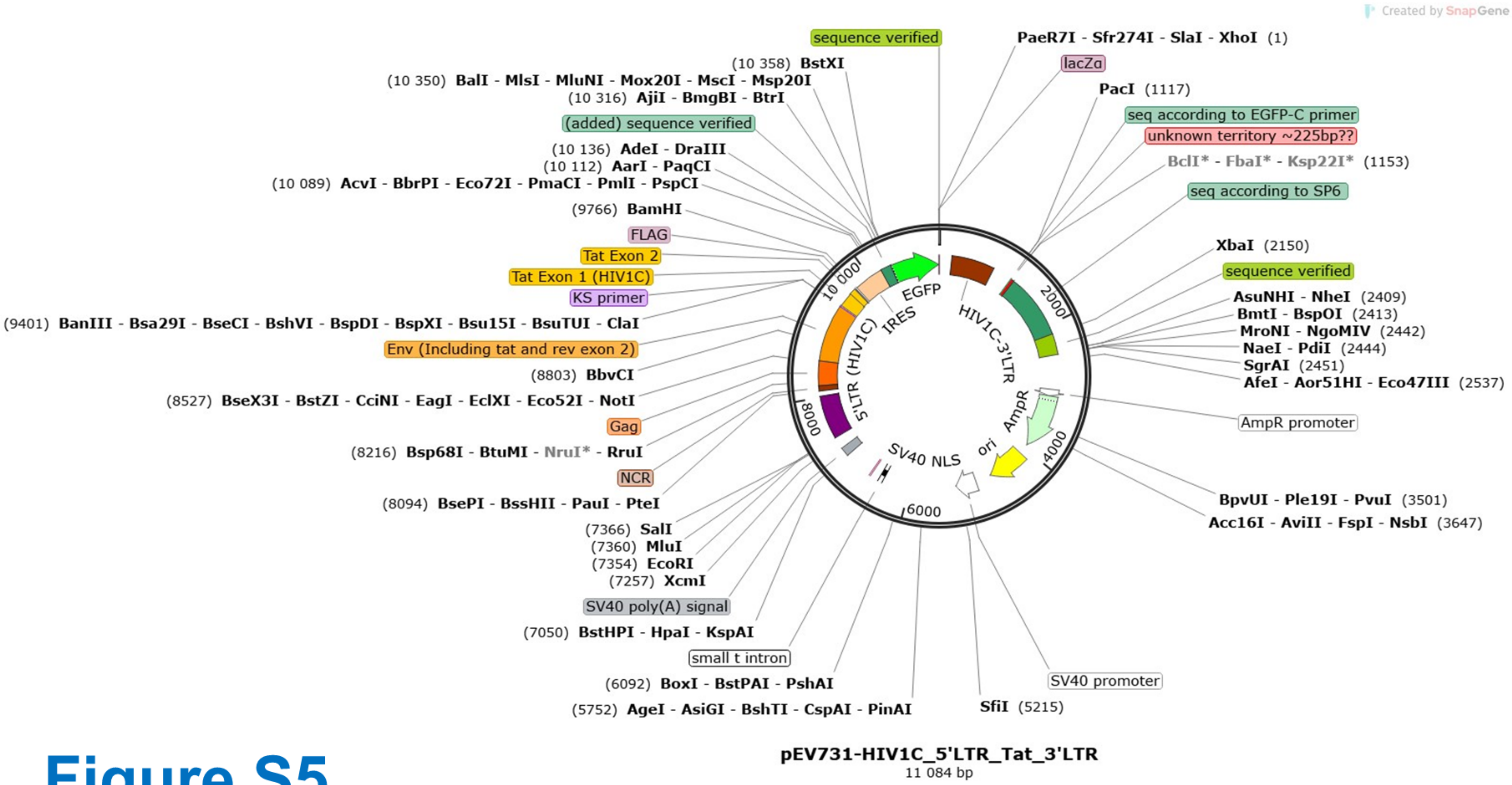
Created by SnapGene
sequence verified
PaeR7I - Sfr274I - SlaI - XhoI (1)
lacZα
PacI (1117)
seq according to EGFP-C primer
unknown territory ~225bp??
BclI* - FbaI* - Ksp22I* (1153)
seq according to SP6
XbaI (2150)
sequence verified
AsuNHI - NheI (2409)
BmtI - BspOI (2413)
MroNI - NgoMIV (2442)
NaeI - PdiI (2444)
SgrAI (2451)
AfeI - Aor51HI - Eco47III (2537)
AmpR promoter
BpvUI - Ple19I - PvuI (3501)
Acc16I - AviII - FspI - NsbI (3647)
SV40 promoter
SfiI (5215)
(5752) AgeI - AsiGI - BshTI - CspAI - PinAI
(6092) BoxI - BstPAI - PshAI
small t intron
(7050) BstHPI - HpaI - KspAI
SV40 poly(A) signal
(7257) XcmI
(7354) EcoRI
(7360) MluI
(7366) SalI
(8094) BsePI - BssHII - PauI - PteI
NCR
(8216) Bsp68I - BtuMI - NruI* - RruI
Gag
(8527) BseX3I - BstZI - CciNI - EagI - EclXI - Eco52I - NotI
(8803) BbvCI
Env (Including tat and rev exon 2)
(9401) BanIII - Bsa29I - BseCI - BshVI - BspDI - BspXI - Bsu15I - BsuTUI - ClaI
KS primer
Tat Exon 1 (HIV1C)
Tat Exon 2
FLAG
(9766) BamHI
(10 089) AcvI - BbrPI - Eco72I - PmaCI - PmlI - PspCI
(10 112) AarI - PaqCI
(10 136) AdeI - DraIII
(added) sequence verified
(10 316) AjiI - BmgBI - BtrI
(10 350) BalI - MlsI - MluNI - Mox20I - MscI - Msp20I
(10 358) BstXI
EGFP
IRES
5'LTR (HIV1C)
HIV1C-3'LTR
AmpR
ori
SV40 NLS
2000
4000
6000
8000
10 000
pEV731-HIV1C_5'LTR_Tat_3'LTR
11 084 bp

### Figure S5

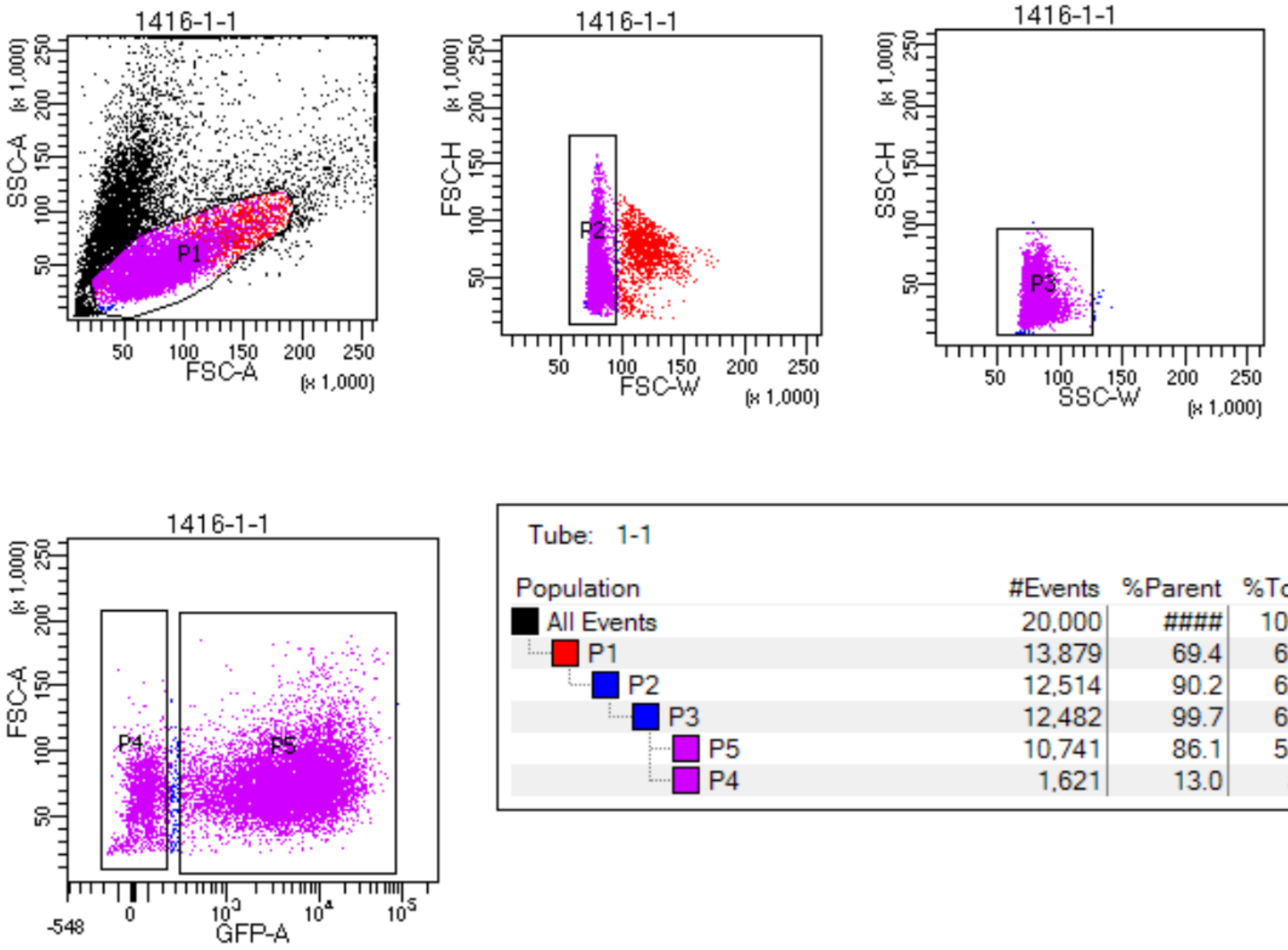


Tube: 1-1

| Population | #Events | %Parent | %Total |
|---|---|---|---|
| All Events | 20,000 | #### | 100.0 |
| P1 | 13,879 | 69.4 | 69.4 |
| P2 | 12,514 | 90.2 | 62.6 |
| P3 | 12,482 | 99.7 | 62.4 |
| P5 | 10,741 | 86.1 | 53.7 |
| P4 | 1,621 | 13.0 | 8.1 |

### Figure S6
